# Genomes of Solenogastres Shed Light on Ancestral Molluscan Biomineralization Genes

**DOI:** 10.64898/2026.11.06.764910

**Authors:** Meghan K. Yap-Chiongco, Kevin M. Kocot

## Abstract

Aplacophora (Solenogastres + Caudofoveata) is a group of worm-like molluscs that lack a shell and instead are covered with aragonitic spines and scales called sclerites. Sclerites are also found along the girdle in chitons (Polyplacophora) and are a synapomorphy for Aculifera (Aplacophora + Polyplacophora), which is the sister taxon of all other shell-bearing molluscs (Conchifera). However, whether the same genetic mechanism underlies the process of biomineralization in aculiferans and conchiferans is unknown. To investigate evolution of genes involved in biomineralization across Mollusca, we re-sequenced the genomes of two species of Solenogastres, *Epimenia babai* and *Neomenia megatrapezata*, producing near-chromosome-level assemblies through PacBio HiFi and Hi-C scaffolding. Using a combination of orthology inference and similarity searches against a known SMP database we identified 1,244 putative SMP orthogroups across Mollusca, 292 of which were present in at least one solenogaster genome. SMPs undergoing significant expansion/contraction at key nodes were further investigated. Of these, 15 were found in at least one solenogaster genome and included families containing chitin-binding, von Willebrand factor A, and C-type lectin domains. Across Mollusca, Solenogastres showed the highest number of significant losses in all gene families, including the SMPs investigated. Targeted searches for structural, silk-like proteins in Aculifera show further reductions in Solenogastres (16-17) compared to caudofoveates (29) and chitons (46-80). Altogether, our results support a conserved but reduced biomineralization toolkit underlying sclerite formation in Solenogastres and identify candidate genes for further proteomic and functional study.

**Article Summary:** While most molluscs (snails, clams, etc.) are known for their iconic shell, aplacophorans (Solenogastres + Caudofoveata) lack this structure and are instead covered by small spines and scales called sclerites. The authors explore the genes involved in sclerite production by sequencing two solenogaster genomes. They conduct comparative analyses across Mollusca and assess the mineralization toolkits of shell- and sclerite-producing molluscs. They show similarities in genes used for both structures, but large losses in solenogasters. Overall the authors provide high-quality genomic resources for Solenogastres and new insights into the evolution of molluscan mineralized structures.

## 1. Introduction

The process by which organisms produce mineralized hard parts is called biomineralization (Marin 2020). This process occurs across the tree of life but is exemplified in Mollusca whose members produce a variety of mineralized structures including the iconic molluscan shell. Mollusca is divided into two major clades, Aculifera (Polyplacophora and Aplacophora) and Conchifera (all remaining classes), a classification well-supported by phylogenomic studies (Kocot et al. 2011; Smith et al. 2011; Vinther et al. 2012; Song et al. 2023; Chen et al. 2025). The scleritome refers to the body of mineralized structures present in a given mollusc, consisting of the shells found in Conchifera, the calcareous scales and spines (sclerites) of Aculifera, and the shell valves of chitons (Polyplacophora)(Checa et al. 2017).

Molluscan shells are organo-mineral biocomposites with the mineral component made of layers of calcium carbonate (CaCO_3_). These structural layers differ in crystallography (generally calcite or aragonite) and their three-dimensional arrangement (Belcher et al. 1996; Checa 2018). CaCO_3_ crystals make up the majority of the shell with only ∼1-5% of total weight being organic matter, termed the shell matrix (Marin 2020). The shell matrix is composed of a complex combination of polysaccharides, lipids, and proteins and differs significantly among molluscan groups (Checa 2018). The shell matrix has been shown to be fundamental in the regulation of shell formation and provides structural toughness (Marin 2020; Huang and Zhang 2022). Most studies on the shell matrix have focused on proteins secreted by the mantle that become trapped within the shell during calcification (Huang and Zhang 2022). These shell matrix proteins (SMPs) have been shown to play critical roles in CaCO_3_ precipitation serving functions such as crystal nucleation, regulation of shell microstructure morphology, and crystal growth/inhibition (Suzuki et al. 2013; Clark 2020; Marin 2020; Yarra et al. 2021; Huang and Zhang 2022). Marin (2020) characterised common domains of SMPs into various categories—extracellular matrix (ECM) or ECM-binding, enzymatic, saccharide-binding, immunity and protection, and cytoplasmic and nuclear proteins. While the domain profiles of SMPs are highly variable, studies have identified domains common among SMPs such as carbonic anhydrase, tyrosinase, chitin-binding, and von Willebrand Factor A (VWA) (Kocot, Aguilera, et al. 2016; McDougall et al. 2016; McDougall and Degnan 2018; Marin 2020). Comparative proteomic and transcriptomic studies have also shown the presence of repetitive low complexity domains (RLCDs) in SMPs found across Conchifera (Jackson et al. 2010; McDougall et al. 2016; Marin 2020; Zhang et al. 2021). Silk is an elastic and strong RLCD-containing protein produced by many metazoans (McDougall et al. 2016). The features characteristic of spider silks are also found in proteins responsible for tough, extracellular structures in a variety of other organisms including the molluscan shell (McDougall et al. 2016). These silk-like shell proteins (SLSPs) include functional domains interspersed with highly repetitive regions with low sequence complexity (McDougall et al. 2016; Jung et al. 2021). Plasticity in SLSP domain profiles, RLCDs, replication slippage, and exon-shuffling contribute to a high level of sequence novelty that is observed in the secretomes between even closely related species (Kocot et al. 2016; Aguilera et al. 2017).

Shells are present in at least some members of all classes within Conchifera and in aculiferan polyplacophorans (chitons). The eight dorsal shell plates of chitons are composed of three layers with cross-lamellar microstructures similar to what is observed in gastropod shells (C. Liu et al. 2024). The configuration of the cuticle with underlying girdle mantle epithelium is proposed to be homologous to the periostracum and associated inner portion of the mantle fold in bivalves (Checa et al. 2017). However, due to differences in structure and developmental origins, the strict homology of chiton shell plates to conchiferan shells remains ambiguous (Furuhashi et al. 2009; Scherholz et al. 2013; C. Liu et al. 2024). Chitons also bear scale-or spine-like sclerites; as the sclerites of chitons grow, neighbouring cells progressively aid in their secretion through the formation of an elevated papilla (Haas et al. 1981; Henry et al. 2004; Checa et al. 2017). While both shells and sclerites are formed by mantle tissue, cell fate mapping has shown that chiton sclerites and shell plates are produced by cells derived from different micromeres during development (Henry et al. 2004), suggesting the two structures are not strictly homologous.

To unravel the evolutionary relationship between chiton sclerites and shell plates to one another and to the conchiferan shell, recent studies have focused on the underlying molecular mechanisms of chiton biomineralization. Whole genome and transcriptome analysis of the chiton *Acanthopleura granulata* by Varney et al. (2021) found the presence of carbonic anhydrase and *Pif-like* genes in the genome and high expression of the Pif97 homolog in the sclerite-secreting girdle tissue. The role of *Pif-like* genes in chiton mineralization was further supported by Lui and colleagues in 2024 who found two SMPs, *Pif-like* and *Nacrein-like*, in the proteome of the shell plates of the chiton *Acanthopleura loochooana* (C. Liu et al. 2024). They expanded upon this study with a proteomic investigation of the sclerites of *A. loochooana* (H. Liu et al. 2024) in which they identified shared and unique sclerite/shell plate SMPs (H. Liu et al. 2024). Shared sclerite/shell plate SMPs included novel glycine-rich proteins in addition to known biomineralization proteins such as carbonic anhydrases, tyrosinase-hemocyanin, cadherin, and von Willebrand factor type A (H. Liu et al. 2024). The authors concluded that, based upon key shared sclerite/shell SMPs, the sclerites and valves of chitons are the result of a shared mineralization toolkit, despite many genes unique to shell plates or sclerites (H. Liu et al. 2024).

While chiton sclerites are restricted to the edge of the girdle, sclerites of the worm-like Aplacophora (Solenogastres + Caudofoveata) cover virtually the entire body of these shell-less animals. Similarly to chitons, aplacophoran sclerites are composed of aragonite (Rieger and Sterrer 1975; Scheltema and Ivanov 2002; Wendt et al. 2023) but are formed by a single cell which invaginates to create a crystallisation chamber in which extracellular formation of the sclerite takes place (Haas et al. 1981; Kingsley et al. 2013; Castro-Claros et al. 2024). Further differences were revealed by recent crystallographic studies of caudofoveate and solenogaster sclerites showing that each individual sclerite is formed from a single crystal of aragonite (Wendt et al. 2023; Castro-Claros et al. 2024). Individual sclerites are separate mineralized entities with limited contact with one another during formation. As such, epithelial cells and the associated gene expression and secreted proteins determine aragonite crystal orientation and thus sclerite texture and morphology (Castro-Claros et al. 2024). This finding demonstrates the important role of secreted biopolymers in solenogaster sclerite formation (Castro-Claros et al. 2024; Castro-Claros et al. 2026), but no studies to date have investigated the molecular basis of aplacophoran sclerite mineralization.

Although molluscan shell biomineralization has been extensively studied, the genomic basis of biomineralization in shell-less aculiferans remains largely unexplored. Due to their phylogenetic position (Kocot et al. 2011; Smith et al. 2011; Vinther et al. 2012; Kocot et al. 2020) and complete lack of shells, Solenogastres represents a key lineage for inferring the ancestral biomineralization toolkit of Mollusca. Although shell matrix proteins (SMPs) can be highly divergent, some protein families and domains associated with biomineralization are conserved across Mollusca. This conservation provides a basis for using comparative genomics to identify candidate biomineralization genes and investigate the evolution of the molluscan biomineralization toolkit. Here, we generated near-chromosome-level genome assemblies of two solenogaster species, *Epimenia babai* Salvini-Plawen, 1997 and *Neomenia megatrapezata* Salvini-Plawen and Paar-Gausch, 2004. Using these genomes and comparative analyses of other molluscan genomes, we investigated the conservation, expansion, and contraction of gene families associated with biomineralization to gain insight into the evolution of molluscan scleritomes.

## 2. Methods

### A. Sampling and Sequencing

The same individual of *Epimenia babai* used for whole genome sequencing by Yap-Chiongco et al. (2024b) was used in this study (see Okusu, 2002 for collection details). The specimen of *Neomenia megatrapezata* (Ap212) used here for genome and Hi-C sequencing was collected on 25 January 2013 near Wrights Gulf in Antarctica (NBP12-10 Station 13, -73.296662, -129.192443 506 m depth) via Blake trawl. This specimen is deposited in the Alabama Museum of Natural History (ALMNH) under catalogue number ALMNH:Inv:25735.

A piece of the midbody of *E. babai* and gonad tissue from *N. megatrapezata* were frozen at -80°C and sent to U.C. Davis DNA Technologies core for high-molecular-weight DNA extraction. Libraries for PacBio sequencing were constructed using the HiFi v2.0 kit for *E. babai* and the HiFi Ultra-Low Input protocol (PacBio, Menlo Park, CA, USA) for *N. megatrapezata*. Using the PacBio Sequel II platform, we sequenced one flow cell for *E. babai* and two flow cells for *N. megatrapezata*.

Hi-C library preparation was performed in-house using the Phase Genomics Hi-C Animal Kit KT2030 v.3 (Phase Genomics, Seattle, WA) with 0.2 g of tissue following manufacturer instructions. Libraries were sequenced on an Illumina NovaSeq S4 flow cell at Psomagen (Cambridge, MA) with 2X150 bp reads.

### B. Genome Assembly, Quality Assessment, and Scaffolding

For *E. babai,* HiFi reads with accuracy above QV 20 (0.99) were extracted using extracthifi (https://github.com/PacificBiosciences/pbtk#extracthifi). HiFi reads from *N. megatrapezata* were adapter trimmed using HifiAdapterFilt v.2.0.0 (Sim et al. 2022) with the options “-l 23” and “-m 95”. Raw Hi-C reads were cleaned and adapter trimmed using matlock with default settings (https://github.com/phasegenomics/matlock). Phased assemblies were produced using filtered HiFi and Hi-C reads as input for hifiasm v.0.16 with the option “-l 3” (Cheng et al. 2021). Assemblies were filtered based on GC content, non-metazoan BLAST hit, and coverage using Blobtools2 (see Blobtools parameters on FigShare) (Challis et al. 2020).

Genome size was estimated using KMC 3 (Kokot et al. 2017) (-k21 -fm -t16 - m500 -ci1 -cs10000) to obtain k-mer statistics followed by kmc-tools with the transform option to create a histogram for input into GenomeScope2 (Ranallo-Benavidez et al. 2020). Genome quality and completeness were assessed after each step involving filtering or polishing of the assembly with QUAST 5.0.2 (Mikheenko et al. 2018) and BUSCO 4.2.0 (Manni et al. 2021) using the Metazoa odb_10 dataset and the “--long” flag.

The post-hifiasm assembly of *N. megatrapezata* and the post-Blobtools2 filtered assembly of *E. babai* were used for scaffolding with Hi-C reads using the Arima Mapping Pipeline (https://github.com/ArimaGenomics/mapping_pipeline). The resulting quality-filtered BAM file was then used as input for scaffolding using YaHS v.1.2a.2 (Zhou et al. 2023). The final output of YaHS was then converted into .hic and .assembly format using juicer tools (Durand et al. 2016) for assessment and manual curation in Juicebox Assembly Tools (JBAT)(Dudchenko et al. 2018).

### C. Repeat Masking and Genome Annotation

For genome annotation, repeats in the final assemblies were annotated and softmasked with RepeatMasker v. 2.0.3 with the “--slow” and “--gccalc” options based on a custom repeat database generated for each species with RepeatModeler v. 2.0.3 (Smit et al. 2015) with the “--LTRStruct” option. The engine used for both programs was rmblast.

Following Yap-Chiongco et al. (Yap-Chiongco et al. 2024), the proteomes of other aplacophorans (See Supplementary Table S1) were mapped to the final genome assemblies with ProtHint 2.6 (Brůna et al. 2020) with an e-value cutoff of 1e-25. Transcriptome reads for each species (NCBI SRX23536359, SRR331899, and SRX23536358 for *N. megatrapezata*; SRR6926338 and SRX23536357 for *E. babai*) were adapter and quality trimmed and filtered using TrimGalore (Krueger et al. 2023) (-q 30 --illumina --trim-n --length 50) and mapped to the respective masked genomes using STAR 2.4.0k (Dobin et al. 2013). Annotation of protein-coding genes was performed with BRAKER 2.1.6 (Brůna et al. 2021) using the output of ProtHint (Brůna et al. 2020) and STAR with the following settings: “--eptmode --softmasking --fitlerOutShort --crf.” Predicted transcripts were retained if they received support from STAR read mapping and/or ProtHint proteome mapping using the selectSupportedSubsets.py bundled with BRAKER.

### D. Hox Gene Annotation

While SMPs involved directly in biomineralization are diverse, highly conserved regulatory genes are also an important part of the molluscan “biomineralization toolkit.” Therefore, the annotation of Hox and other homeobox genes was performed following Leite et al. (Leite et al. 2022). Briefly, hmmbuild v.3.3 (Eddy 2011) was used to create an HMM using the homeodomain PF0046 Pfam RP55 alignment to query protein annotations with hmmsearch v.3.3 (Eddy 2011). Domain hits were extracted with elsfetch v. 0.49, retaining hits >50 amino acids in length for further investigation. To increase the representation of molluscan species the same approach was used to extract Hox genes from publicly available genome annotations for two scaphopods, two chitons, and one caudofovete. Other molluscan Hox genes were downloaded from NCBI (Supplementary Table S2) and aligned with hits using MAFFT-DASH (Rozewicki et al. 2019). Alignments were refined and manually checked in AliView v.1.27 (Larsson 2014). The final alignment was then used for phylogenetic inference using maximum likelihood in IQ-Tree 2.1.3 (Minh et al. 2020) using the best-fitting model (-m MFP) with 1000 rapid bootstraps.

### E. Orthology Inference and Phylogenetic Reconstruction

Protein annotations from high-quality, publicly available genomes of 29 molluscs, two annelids, one nemertean, and three lophophorates were used along with the protein annotations of *N. megatrapezata* and *E. babai* for orthology inference using OrthoFinder 2.4.0 (Emms and Kelly 2019). Matrix refinement followed the approach of Yap-Chiongco et al. (Yap-Chiongco et al. 2024) using ClipKIT 2.2.6 for trimming of ambiguously aligned regions and retaining only genes sampled from at least 18/34 taxa. The concatenated supermatrix was used for maximum likelihood phylogenetic analysis with IQ-Tree 2.1.3 (Minh et al. 2020) using the best-fitting model for each partition (-m MFP) with 1000 rapid bootstraps.

### F. Gene Family Analyses and Identification of Potential SMPs

CAFE5 (Mendes et al. 2021) was used on the output of OrthoFinder to infer gene gain and loss over time. CAFE5 employs a stochastic birth-death probabilistic model to estimate the rate of change across a given tree. The program calculates a p-value for each gene family to determine whether it has a significantly greater rate of evolution than average and then pinpoints the nodes in which these significant changes occurred using a Viterbi p-value (Mendes et al. 2021). The Orthogroups.GeneCount.tsv file was reformatted for use as input to CAFE5 and filtered using the CAFE5 clade_and_size_filter.py script to exclude gene families with high gene copy number variance. After filtering, a total of 21,854 orthogroups were retained for analysis. InterProScan 5.360-92.0 (Jones et al. 2014) was then used with the “-iprlookup - goterms -appl Pfam -b” settings to functionally annotate each orthogroup. An ultrametric tree was obtained from the tree produced by IQ-Tree using R8s v.1.81 (Sanderson 2003). The configuration file for R8s was obtained by modifying the output of the CAFE5 script prep_r8s.py to include fossil calibration points following the priors outlined by Song et al. (Song et al. 2023). CAFE5 was first run to estimate an error file with the flags “-e” and “-p.” The resulting error file was then used with “-e” for error correction in subsequent analyses using CAFE5. CafePlotter (https://github.com/moshi4/CafePlotter) was used to visualise expansion/contraction of gene families on the phylogeny (Figure 1).

**Figure 1.**
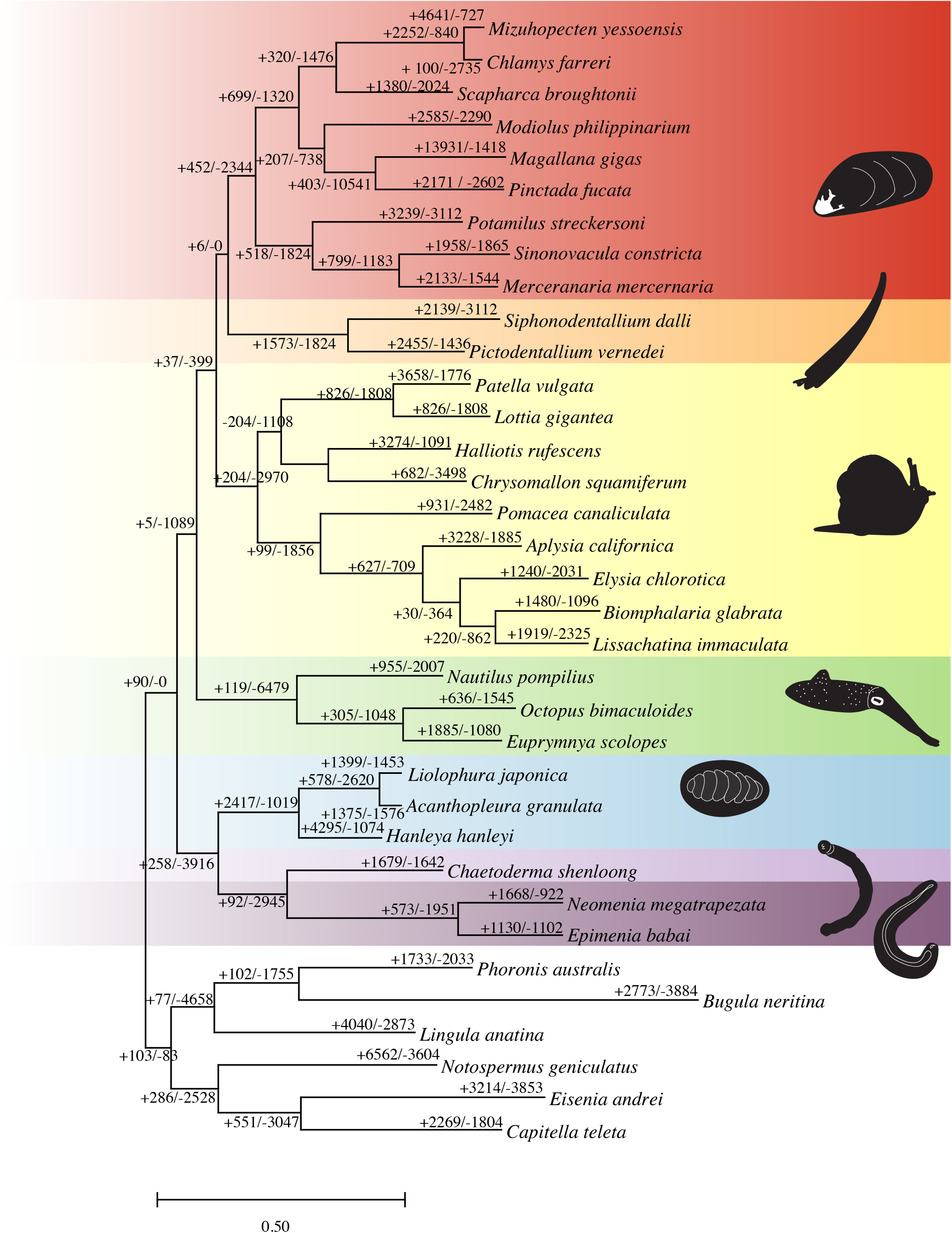
Phylogenetic reconstruction of Mollusca based on whole genomes. Results of CAFE5 analysis of gene family expansions (+) and contractions (–) are indicated along each branch. Colors and accompanying silhouettes denote molluscan classes. Scale bar indicates substitutions per site.

To study gains and losses of potential biomineralization genes within Aculifera, orthogroups were filtered to identify 3,291 likely involved in biomineralization based on functional annotation of domains and GO terms. BLASTP (Camacho et al. 2009) was used with an e-value of 1e-12 to search gene models against a curated molluscan SMP database (https://doi.org/10/cz2w; Yarra 2019) that includes 327 molluscan SMPs. As the SMP Database is relatively small and biased towards conchiferan taxa, percent identity and alignment length were considered in assessing the output, with a putative SMP only included if it had an alignment length over 100 and percent identity >25%.

### G. Identification of Silk-Like Structural Proteins

To infer silk-like structural proteins (SLSPs), we implemented the steps of SilkSlider (McDougall et al. 2016) using a custom python script (https://github.com/mkyapchiongco/Modified-SilkSlider). SLSPs are characterised by the presence of a signal peptide and a glycine rich domain (McDougall et al. 2016). First, glycine-rich sequences were identified using the same approach as McDougall et al. 2016. As SLSPs are trafficked outside of the cell membrane, the exclusion of sequences without a signal peptide ensures the exclusion of any extracellular debris (McDougall et al. 2016). Signal-P v. 5 (Almagro Armenteros et al. 2019) was then used to exclude any sequences lacking a signal peptide followed by DeepTMHMM (Hallgren et al. 2022) to exclude sequences with a transmembrane domain. InterProScan5 (https://www.ebi.ac.uk/jdispatcher/pfa/iprscan5) was used for domain annotation of the final silk-like proteins. To assess similarity to known SLSPs, predicted sequences were searched against the NCBI nr database using blastp (Camacho et al. 2009).

## 3. Results

### A. Sequencing

Sequencing of two PacBio HiFi SMRT cells for *N. megatrapezata* yielded a total of 5.34 M reads. Only 93 reads were excluded by HifiAdapterFilt. For *E. babai*, extracthifi was used to retain high-quality reads from one flow cell of HiFi sequencing resulting in 1.45 M reads above QV 20.

### B. Genome Assembly, Quality Assessment, and Scaffolding

GenomeScope analysis based on HiFi reads estimated the genome size of *N. megatrapezata* to be 272 Mbp with a heterozygosity of 1.53%. Assembly with hifiasm yielded an assembly of 4022 contigs (N50 = 0.339 Mbp, L50 = 448) with 93.9% complete BUSCOs detected (7.8% duplicated, 2.2% fragmented, and 3.9% missing). After removal of contaminants and scaffolding, the final *N. megatrapezata* assembly was reduced to 2713 contigs (N50 = 13.47 Mbp, L50 = 14) with a BUSCO completeness score of 93.9% (6.4% duplicated, 2.0% fragmented, and 4.1% missing). The final *N. megatrapezata* assembly is near chromosome level with 72.34% of reads mapping to 24 primary scaffolds (Supplementary Figures S1-2 and Tables S3-S5).

For *E. babai*, GenomeScope estimated a genome size of 442 Mbp with a heterozygosity of 2.25%. Hifiasm yielded an assembly of 2545 contigs (N50 = 4.977 Mbp, L50 = 40) with a BUSCO completeness score of 87.9% (6.5% duplicated, 2.4% fragmented, and 9.7% missing). After removal of contaminants and Hi-C scaffolding, the final *E. babai* assembly was reduced to 520 contigs (N50 = 17.336 Mbp, L50 = 11) with a BUSCO completeness score of 93.0% (0.9% duplicated, 1.6% fragmented, and 5.4% missing). The final assembly is near chromosome level with 88.32% of reads mapping to 23 primary scaffolds.

### C. Repeat Masking and Genome Annotation

Repeat masking of the *N. megatrapezata* genome resulted in 45.30% of bases masked, largely composed of unclassified repeats (23.96%) and LINE retroelements (14.82%). Annotation using BRAKER2 resulted in 20,095 gene models with a BUSCO completeness score of 95.1% (10.7% duplicated, 2.2% fragmented, and 2.7% missing). Of these, 16,896 gene models were supported by RNAseq and proteins with this subset receiving a BUSCO completeness score of 95.2% (10.7% duplicated, 2.3% fragmented, and 2.5% missing). Functional annotation resulted in the identification of at least one GO term in 8,141 genes and 13,154 genes with one or more PFAM match (4,954 unique PFAM domains). Of the genes supported by functional annotation, transcript, and/or protein evidence, 76.9% map to one of the 24 primary scaffolds.

For *E. babai*, repeat masking resulted in 44.85% of bases masked, which were mainly identified as unclassified repeats (24.02%). Annotation using BRAKER2 resulted in 22,264 gene models with a BUSCO completeness score of 91.4% (4.7% duplicated, 4.0% fragmented, 4.6% missing). Of these, 15,268 were supported by RNAseq and proteins with this subset of the gene models having a BUSCO completeness score of 91.2% (4.7% duplicated, 3.9% fragmented, 4.9% missing). Of the supported gene models, at least one GO term was identified for 7,449 genes, a PFAM match was found for 11,821 genes (a total of 4,873 unique PFAM domains), and 97.9% of supported genes map to one of the 23 primary scaffolds.

### D. Orthology Inference and Phylogenetic Reconstruction

Our phylogenomic pipeline resulted in a final concatenated matrix of 6,555 orthogroups totaling 167,204 amino acid positions with 41.50% missing data. Relationships within Mollusca (Figure 1) were strongly supported and in agreement with the accepted dichotomy of Conchifera + Aculifera. Notably, we recovered Scaphopoda as the sister taxon to Bivalvia (Diasoma) in accordance with other recent studies (Song et al. 2023; Chen et al. 2025).

### E. Gene Family Analyses and Identification of Potential SMPs

Of 21,854 orthogroups, CAFE5 identified a total of 1,113 orthogroups as having undergone significant expansions and/or contractions (Figure 1). For Conchifera, we observed significant expansion of only five families and significant contraction of only one while, in contrast, we observed 58 expansions and six contractions at the base of Aculifera. Across Aculifera, most significantly changed gene families were expanded, with reduction happening primarily along the Solenogastres branch. In some cases, gene families were expanded within Polyplacophora and Caudofoveata but contracted within Solenogastres (Figure 1). For Aplacophora, eight gene families were significantly expanded and 55 gene families were significantly contracted. There is another large contraction of 115 gene families for Solenogastres, seven of which contain domains involved in biomineralization (chitin-binding, von Willebrand factor type A, *Pif*, and C-type lectin).

A total of 1,244 orthogroups were identified as putative molluscan SMPs 292 of which were present in at least one solenogaster genome. Of the total putative SMPs, 74 were significantly expanded or contracted over the tree with 55 of these present in at least one solenogaster genome. Candidates were further filtered to include only putative SMPs that have undergone significant expansion or contraction at Mollusca, Conchifera, Aculifera, Polyplacophora, Aplacophora, Caudofoveata, or Solenogastres nodes, resulting in a final set of 21 orthogroups (Supplementary Table S6). These 21 orthogroups include hits to perlucin, tyrosinase, ependymin, EGF-like, fibronectin-containing, beta-hexosaminidase, and chitin synthase.

Seven orthogroups were identified as the nacreous layer forming SMPs *Pif-like* or *blue mussel shell protein (BMSP)-like* (Supplementary Table S7) and show dynamic expansion and contraction within Mollusca. OG0000057 on average had the highest percent identity and longest alignments to *Pif* sequences in the SMP database curated by Yarra (2019). We see a significant expansion of OG0000057 at the bases of Mollusca and Aculifera. This gene family is expanded at the base of Aplacophora, at the base of Solenogastres, and in the chiton *Hanelya hanleyi*. In conchiferan molluscs, there are expansions in Patellogastropoda, particularly in the patellogastropod *Patella vulgata,* and in the scaphopod *Pictodentallium vernedei*. A contrasting pattern was observed in OG0000363, which underwent an inferred expansion at the base of Aculifera but was subsequently lost in all sampled aplacophoran lineages. CAFE5 analysis indicates a significant contraction from three copies to one at the base of Aplacophora, followed by further reductions to zero copies in the sampled lineages. OG0000363 was also absent from the sampled pulmonate gastropods but was retained in other sampled molluscan lineages, including chitons.

### F. Identification Silk-Like Structural Proteins

We identified 17 silk-like structural proteins (SLSPs) in the genomes of *E. babai* and 16 in *N. megatrapezata* (Supplementary Table S8). For comparison, 29 SLSPs were identified in the caudofoveate aplacophoran *Chaetoderma shenloong*, 46 in *Liolophura japonica,* 57 in *Acanthopleura granulata*, and 80 in *Hanleya hanleyi*.

Nine of the 17 identified SLSPs of *E. babai* had BLAST hits to the molluscan SMP database whereas only four of the 16 SLSPs predicted from *N. megatrapezata* had a positive hit. Six of these SLSPs (five found in *Epimenia*, one found in *Neomenia*) belong to one orthogroup (OG0000065) and blast to tyrosinase domains of SMPs originally isolated from bivalves (Supplementary Table S8).

### G. Hox Gene Annotation

Both *N. megatrapezata* and *E. babai* have all 11 molluscan Hox genes (Figure 2; Supplementary Figure S3). The Hox cluster of *N. megatrapezata* was found primarily on one scaffold (Scaffold 33) with only *Hox3* falling outside of this cluster (Scaffold 63). Further we observed a deviation from the inferred ancestral molluscan Hox gene order, with posterior Hox genes *Post1* and *Post2* upstream of *Hox1* by 0.14 Mbp. The ten Hox genes on Scaffold 33 form a cluster 0.66 Mbp long.

**Figure 2.**
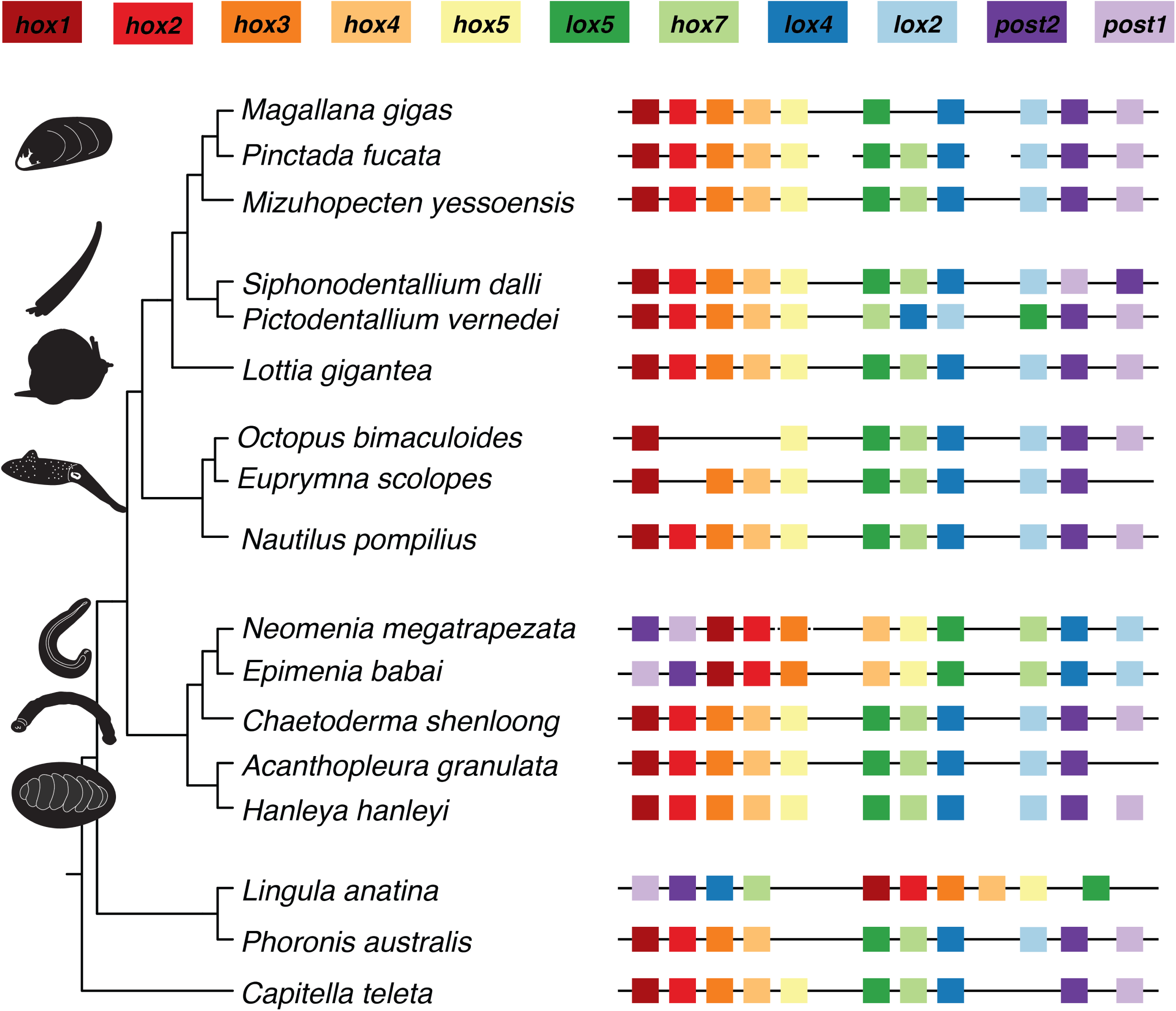
Synteny of Hox genes in Solenogastres and other taxa. The presence of a gene is noted by a box of the corresponding color. If the cluster was recovered on one contiguous scaffold, this is noted by a continuous line. Broken lines represent recovery of the cluster on more than one scaffold. The dashed arrows represent a possible inversion in Solenogastres of *Post1* and *Post2* genes.

In *E. babai*, these genes were all recovered on one scaffold (Scaffold 17). *Hox1* through *Lox2* are within a 0.45 Mbp cluster, while the posterior Hox genes *Post1* and *Post2* are upstream of *Hox1* by 6.01 Mbp.

## 4. Discussion

### A. High-Quality Genomes of Two Solenogastres

Yap-Chiongco et al. (2024) sequenced draft genomes for both *N. megatrapezata* and *E. babai* using a hybrid assembly approach with Oxford Nanopore long reads and Illumina short reads. Compared to those draft genome assemblies, the HiFi-based genomes presented in this work are more contiguous and both the assemblies and gene models have higher BUSCO scores (Supplementary Table S2). *E. babai* and *N. megatrapezata* assemble into 23 and 24 pseudomolecules, respectively, with estimated sizes of 442 Mbp for *E. babai* and 272 Mbp for *N. megatrapezata*.

While not all scaffolds were able to be placed within Hi-C pseudomolecules, these genomes are comparable in terms of BUSCO score and scaffold N50 to chromosome-level molluscan genome assemblies (see Goodheart et al. 2024) and Supplementary Figure S2). The *E. babai* and *N. megatrapezata* assemblies contain a relatively high level of repeats (44.85% and 45.30%, respectively) and high heterozygosity (2.25% and 1.53%, respectively), similar to what is known from other molluscs (Gomes-dos-Santos et al. 2020; Goodheart et al. 2024).

Yap-Chiongco et al. (2024) noted the differences in size estimates between Oxford Nanopore-based assemblies and Feuglen image analysis densitometry (FIAD) by Kocot, Jeffery, et al. (Kocot, Jeffery, et al. 2016). Here we see further deviation from the FIAD-predicted 750 Mbp genome size for *E. babai* but a closer assembly size for *N. megatrapezata* to the predicted 293 Mbp genome size of the closely related species *Neomenia permagna* (Kocot, Jeffery, et al. 2016). Molluscan genome sizes are diverse (Kocot, Jeffery, et al. 2016; Adachi et al. 2021) and can vary drastically even within members of the same genus (Sigwart et al. 2021). Chitons have variable genome sizes (0.66-2.52 Gbp) (Kocot, Jeffery, et al. 2016) while the genome of *Chaetoderma shenloong* predicted at 2.45 Gbp both estimation via FAID and sequencing (Wang et al. 2024). Reduction in genome size is often associated with changes in gene number, genome architecture, and repeat content (Fernández and Gabaldón 2020; Martín-Durán et al. 2021; Liao et al. 2023). Having comparable levels of repeat content and lower number of gene models in comparison to other molluscan genomes, genome reduction in Solenogastres may be due to loss of genes rather than repetitive content. This is further supported by the large number of contracted gene families observed for the clade in our CAFE5 analysis, but requires further investigation into other mechanisms of potential reduction.

While it is tempting to associate the morphological simplicity of the solenogaster body plan with small genome size, the large genome size of the similarly simplified *C. shenloong* supports the idea that genome size is not related to organismal complexity (Elliott and Gregory 2015; Martín-Durán et al. 2021) and that these associations are likely due to lineage-specific adaptations and the cumulative effects of diverse evolutionary processes.

### B. Biomineralization Genes

Functional annotation and orthology inference revealed the presence of genes known to be involved in conchiferan biomineralization within the genomes of *E. babai* and *N. megatrapezata*. Fifteen of the 21 investigated biomineralization orthogroups were found in at least one solenogaster genome. These genes have domains that have been suggested to be part of an ancient bilaterian biomineralization toolkit, with domains such as chitin-binding, von Willebrand Factor A (VWA), and C-type lectin (McDougall and Degnan 2018; Marin 2020; Zhang et al. 2021). While the presence of these genes across Mollusca suggests conservation, we observed low sequence similarity within biomineralization orthogroups. Previous studies have shown the rapidly evolving nature of SMPs even among closely related species (Aguilera et al. 2014; Aguilera et al. 2017; Zhang et al. 2021).

Of the 21 SMP orthogroups investigated, Solenogastres show the greatest loss in comparison with other molluscs. This is in line with the overall trend of expansion and contraction of gene families where Solenogastres also showed the highest number of contractions. A potential reduction in genome size with respect to other aculiferans may have contributed to the observed losses of biomineralization genes. The reduced solenogaster SMP toolkit may help to identify what domains are minimally necessary for biomineralization. Many SMPs are known to serve multiple functions during biomineralization and are able to regulate crystal nucleation and growth and induce changes in crystal morphology (Marin et al. 2007; Yan et al. 2007; Li et al. 2025). The multifunctionality of some SMPs would allow for the reduction in the biomineralization toolkit of Solenogastres without losing the ability for biomineralization. The expansion and contraction of biomineralization genes shown by our data supports the flexibility of the SMP repertoire across Mollusca as well as the conservation of some key domains (Kocot, Aguilera, et al. 2016; McDougall and Degnan 2018; Marin 2020; C. Liu et al. 2024). While these SMPs have been shown to be involved in shell development in other molluscs, targeted transcriptome sequencing of the sclerite-secreting epithelium in developing larvae and proteomics of the sclerites themselves need to be undertaken to explore the SMPs directly involved in biomineralization in Solenogastres. Further, these results are limited by the small number of solenogaster genomes in the present study. As new solenogaster genomes become available, our results serve as a starting point to further explore these patterns.

### C. Silk-like Proteins

SLSPs found within molluscan shells are known to aid in formation and reconstruction of shell microstructures providing the matrix with elasticity and strength (Jung et al. 2021). The elasticity of these proteins is due to the highly repetitive sequences of amino acids that fold into flexible secondary structures such as β-pleated sheets (Jung et al. 2021). Varney *et al*. (2021) analysed the SLSPs of the chiton *Acanthopleura granulata,* finding 31 putative SLSPs, 23 of which had significant similarity to genes demonstrated to be involved in mineralization in other molluscs. Further, the authors also noted that several silk-like proteins were highly expressed in the girdle of the chiton *A. granulata* (Varney et al. 2021). Our analysis of silk-like proteins encoded in the genomes of five diverse aculiferans (two solenogasters, three chitons, and one caudofoveate) predicted a greater number of SLSPs within chitons as compared to the aplacophorans. While the majority of SLSPs in *A. granulata* predicted by Varney et al. (2021) had significant hits to known mineralization genes, just 12 solenogaster SLSPs had significant hits to known biomineralization genes. These 12 SLSPs fell within seven orthogroups, two of which underwent significant expansion/contraction in Mollusca OG0000027 (proteinase inhibitor I2 and pancreatic trypsin inhibitor Kunitz domain) and OG0000065 (tyrosinase).

### D. Pif/BMSP-like genes

We identified seven orthogroups as *Pif/BMSP-like.* Both *Pif* and *BMSP* are calcium-binding proteins that have roles in crystal formation of aragonite (Suzuki et al. 2013; Shimizu et al. 2022; Shimizu et al. 2024 June 12). Both proteins are ordered from N-terminal to C-terminal, signal peptides, VWA domain, chitin-binding domain, and partial laminin G domains; however *Pif* only contains one VWA domain while *BMSP* contains four (Suzuki et al. 2011; Suzuki et al. 2013; Shimizu et al. 2022; Shimizu et al. 2024 June 12). Shimizu et al. (2024) classified the *Pif/BMSP* genes into eight sub-families with unique sequence features, expression, and taxonomic distribution (Shimizu et al. 2024 June 12). However, this analysis was focused on conchiferan taxa. Transcripts encoding *Pif/BMSP-like* proteins were shown to be expressed in the sclerite secreting girdle of the chiton *A. granulata* by Varney et al. (2021) and presence of these proteins within valve and sclerite proteomes of *A. loochooana* in a recent proteomic study by Liu et al. (2024). Three of our seven *Pif/BMSP-like* families were significantly expanded in Aculifera. Previous work showing the importance of VWA and chitin-binding domains in mineralization along with our results showing the dynamic expansion/contraction in both major molluscan clades suggest a long standing role of VWA and chitin-binding domains in the ancestral molluscan biomineralization tool-kit.

Similarly to what was observed in other gene families (regardless of putative biomineralization role), Solenogastres have had a greater loss in *Pif/BMSP-like* gene families in comparison to other aculiferans. OG0000057 is the only *Pif/BMSP-like* gene that is expanded in Solenogastres. Conserved VWA and chitin-binding domains suggest a role in aragonite mineralization. Expansion of this gene family at the bases of Mollusca, Aculifera, and Aplacophora warrant further investigation into its putative role in aculiferan biomineralization through functional or proteomic study (Supplementary Table 7).

### E. Orthology Inference is Important to Understanding SMP Evolution

BLAST searches against public databases and within the SMP database curated by Yarra (2019) can often show low identity with short alignments (Batzel et al. 2022). As many SMPs have large RLCDs, sequence similarity is generally highest in conserved protein domains, resulting in short alignment length in BLAST results (Batzel et al. 2022). As we chose arbitrary cutoffs for alignment length and sequence identity, we may have missed SMPs with high levels of sequence divergence between taxonomic groups. Authors have previously highlighted the need for orthology inference and phylogenetic analysis of gene families to determine the level of conservation in molluscan SMPs (Batzel et al. 2022; Lopez-Anido et al. 2024). Our results further demonstrate that the rapidly evolving nature of SMPs result in low alignment length and identity to publicly available databases but through orthology inference, deeper levels of conservation can be observed. Orthology inference will be a useful aid in the next steps of uncovering biomineralization genes in Solenogastres using more direct sequencing approaches (i.e. proteomic sequencing and transcriptome sequencing of sclerite secreting cells).

### F. Hox Gene Analysis

Both *E. babai* and *N. megatrapezata* have a complete complement of Hox genes. All Hox genes are on one scaffold in *E. babai* and, in *N. megatrapezata*, all Hox genes except *Hox3* are located on a single scaffold (Figure 2). The scaffold containing *Hox3*, however, is small (48.8 Kbp) and falls within a region of limited Hi-C contacts. The scaffold containing the remainder of the Hox cluster (Scaffold 33) itself includes a region of sparse Hi-C contacts, which may reflect uncollapsed heterozygosity or high repeat content as 44% of the scaffold was masked by RepeatMasker. Therefore, we cannot conclusively determine the lack of a continuous Hox cluster in *N. megatrapezata*. Further, in both *E. babai* and *N. megatrapezata*, *Post1* and *Post2* were found upstream of *Hox1* (Figure 2). While this peculiarity could also be due to limited Hi-C contacts impeding accurate scaffolding, there is precedent. In the annelids *Owenia fusiformis* and *Parescarpia echinospicata*, *Post1* is located upstream of the anterior Hox genes. In oligochaete annelids, *Post1* is often found on a separate scaffold and it is absent in leeches, nemerteans, flatworms, bryozoans, and phoronids (Barucca et al. 2016; Schultz et al. 2024).

It has been inferred that the last common ancestor of Mollusca had an intact Hox cluster with staggered expression (i.e. in a spatial order) along the anterior-posterior axis (Fritsch et al. 2015; Wanninger and Wollesen 2019). Traces of staggered expression have been observed in chitons, scaphopods, gastropods, and cephalopods (Fritsch et al. 2015; Wollesen et al. 2018; Barrera Grijalba et al. 2023). Spatial and temporal colinearity of Hox genes in the last common ancestor of Polyplacophora are supported by an intact cluster in the genome the chiton *Acanthopleura granulata* and observations of staggered expression of Hox genes in chitons (Fritsch et al. 2015; Fritsch et al. 2016; Varney et al. 2021). Along with the intact cluster of *Chaetoderma shenloong* found by Wang et al. (2024), the intact cluster observed in *E. babai* and a mostly(?) intact cluster in *N. megatrapezata* supports the notion that the ancestral aplacophoran likely had a full Hox cluster with subsequent translocation of the *Post1* and *Post2* genes in Solenogastres. This data suggests that the last common ancestor of Aculifera likely showed staggered and potentially co-linear expression of Hox genes, however expression data are needed from both aplacophoran lineages to confirm this assumption.

Hox and parahox genes play an important role in molluscan shell and sclerite formation (Wollesen and Wanninger 2023; Barrera Grijalba et al.). While the shell-forming tissue of brachiopods only show expression of *Hox1*, many conchiferan and aculiferan molluscs express additional Hox genes in the shell/sclerite field (Schiemann et al. 2017; Barrera Grijalba et al.). *Hox1* is expressed in shell field of conciferan molluscs, within the field of shell-plate one in the chitons *Acanthopleura fascicularis* and *Acanthochitona rubrolineata*, and in the sclerite-bearing cells of the solenogaster *Wirenia argentea* (Barrera Grijalba et al.). While conchiferans express *Hox1* across the entire anterior-posterior axis, its expression is restricted to the anterior regions of the mineralizing tissue in aculiferans (Fritsch et al. 2015; Fritsch et al. 2016; Barrera Grijalba et al.). This is theorized to be reflective of the ancestral staggered expression along the anterior-posterior axis of molluscs (Barrera Grijalba et al.).

## 5. Conclusions

Although research on solenogaster biodiversity has steadily advanced, genomic resources remain limited, and the functional biology of these animals is still poorly understood. Here, we have improved solenogaster genome resources through the re-sequencing and assembly of two distantly related solenogasters—*Epimenia babai* and *Neomenia megatrapezata.* While issues with high levels of heterozygosity and repeat content resulted in unplaced scaffolds, both assemblies are near chromosome-level. In comparison to other molluscan genomes, solenogaster genomes are small in size and gene number. Smaller genome sizes in Solenogastres may be due to gene loss rather than purging of TE’s as repeat-content is on par with other molluscan genomes. As one of the last remaining classes without high-quality genomic data, these solenogaster genomes broaden the avenue for understanding themes in molluscan evolution. For example, we were able to identify full Hox clusters with unique lineage-specific translation of *Post1* and *Post2* upstream of the anterior Hox genes.

Using these genomes, we conducted comparative genomic analyses to identify candidate biomineralization genes and investigate gene family evolution across Mollusca, specifically focusing on gene families with at least some members known to be involved in biomineralization. In accordance with the overall lower number of genes in solenogaster genomes compared to most other mollusc genomes, we see a large number of losses at the base of Solenogastres. We were able to identify both known biomineralization genes and novel genes encoding proteins with combinations of domains involved in biomineralization in other molluscs within solenogaster genomes, suggesting some conservation in molecular mechanisms underlying mineralization in even these shell-less molluscs. However, with lineage-specific co-option and rapidly-evolving sequences, functional studies should be undertaken to evaluate the roles of these genes in Aplacophora as a whole. More targeted approaches, including proteomic analyses of sclerites, transcriptomics of sclerite-secreting cells, and investigations of developmental gene expression and function, are needed to better understand the conservation and divergence of biomineralization mechanisms across Mollusca. Proteomic analyses would be a particularly valuable next step, although the small body size of many solenogasters and the difficulty of obtaining sufficient numbers of specimens present practical challenges. Despite these challenges, such approaches will be important for determining the roles of candidate genes in sclerite formation. Our comparative genomic analyses provide insight into the evolution of biomineralization-associated gene families and identify candidate proteins for future functional studies of sclerite formation in Solenogastres.

## Data Accessibility

Raw genomic data for *Neomenia megatrapezata* are available from NCBI under BioProject PRJNA1186338. The *Neomenia* Illumina Hi-C reads are available under accession SRR31359800 and the PacBio HiFi reads are available under accessions SRR31359801-SRR31359802. Raw genomic data for *Epimenia babai* are available from NCBI under BioProject PRJNA1186338. The *Epimenia* Illumina Hi-C reads are available under accession SRR31364375 and the PacBio HiFi reads are available under accession SRR31364374. Notably, the *Epimenia* specimen from which these data were generated is the same individual that was used to generate data under all of the following BioProjects: SAMN08774992, SAMN39711720, SAMN17155388, SAMN11109285, and SAMN39711717. Code is available at: https://github.com/mkyapchiongco/Solenogaster-Genome-Project and https://github.com/mkyapchiongco/Modified-SilkSlider. Extended data are available through FigShare DOI:10.6084/m9.figshare.25119095

## Ethics

This work did not require ethical approval from a human subject or animal welfare committee.

## Declaration of AI Use

No generative artificial intelligence (AI) tools were used in the writing of this manuscript. The figures, analyses, interpretations, and text presented here were produced by the authors.

## Authors’ Contributions

**M.Y.C:** Conceptualization, Data curation, Formal analysis, Investigation, Methodology, Writing – original draft, Writing – review & editing, Resources, Visualization. **K.M.K:** Conceptualization, Funding acquisition, Methodology, Resources, Supervision, Writing – review & editing, Investigation.

## Acknowledgments

We thank Dr. Ken Halanych and the crew and scientists of the Icy Inverts cruises aboard R/V Lawrence M. Gould and Nathaniel B. Palmer for help with the collection of *Neomenia megatrapezata* and Akiko Okusu for sharing samples of *Epimenia babai*. We also thank Drs. Nickellaus G. Roberts, M. Carmen Cobo Llovo, Andreas Hejnol, Michael McKain, Jane Rasco, and John Yoder for helpful comments on earlier stages of the manuscript. Computational work was performed on the University of Alabama High Performance Computing (UAHPC) cluster.

